# The Drosophila FET orthologue Cabeza is an essential cofactor for ETV4-mediated activation of GGAA microsatellite neoenhancers

**DOI:** 10.64898/2026.07.06.736845

**Authors:** Cristina Molnar, Jose Reina, Jaume Mora, Cayetano Gonzalez

## Abstract

The conversion of transcriptionally silent GGAA microsatellites (GGAAµSats) into functional enhancers by FET::ETS oncogenic fusions is a hallmark of Ewing sarcoma. However, emerging evidence implicates non-fused, full-length oncogenic ETS transcription factors in activating these repeats in other malignancies. Evaluating the *in vivo* transcriptional requirements of various human ETS factors in Drosophila, we found that human ETV4 uniquely binds and robustly activates GGAAµSats in a tissue-specific manner. This activation is strongly inhibited by the human ETS repressor ETV6. Taking advantage of low genetic redundancy in Drosophila, we identified Cabeza (Caz), the single fly FET orthologue, as a necessary cofactor for ETV4-mediated transcription at GGAAµSats. Conversely, EWS::FLI1-mediated transcriptional activation of GGAAµSats is entirely independent of endogenous Caz, highlighting the distinct mechanics of covalent tethering versus non-covalent physical complexes. Collectively, our findings provide definitive *in vivo* evidence that non-fused ETS factors cooperate with endogenous FET proteins to drive transcription from silent GGAA repeats, mechanistically validating this regulatory transformation known to operate as an oncogenic mechanism beyond Ewing sarcoma.

## Introduction

The conversion of transcriptionally silent GGAA microsatellites (GGAAµSats) into functional enhancers is widely recognized as a hallmark of Ewing sarcoma. This process is driven by oncogenes composed of the N-terminal domain of a FET family protein (FUS, EWSR1, or TAF15) joined to the DNA-binding domain of an ETS factor such as FLI1, ERG and others. The most frequent of such oncogenes is EWS::FLI1, which is the driver for more than 80% of all Ewing sarcomas (Delattre et al. 1994; Sankar and Lessnick 2011; Crompton et al. 2014; Grünewald et al. 2018). The activation of GGAA microsatellites by these fusion proteins induces a tumor-specific gene regulatory program representing a critical mechanism in the pathogenesis of Ewing sarcoma (Gangwal et al. 2008; Guillon et al. 2009; Patel et al. 2012; Johnson et al. 2017; Boulay et al. 2018; Riggi et al. 2021).

ETS proteins play critical roles in normal tissue development and homeostasis and have been implicated as key drivers in a variety of human cancers. ETS proteins are characterized by a highly conserved 5’-GGA(A/T)-3’ DNA binding motifb (Sharrocks et al. 1997; Hsu et al. 2004; Hollenhorst et al. 2011; Cooper et al. 2015). Although many ETSs share the ability to bind GGAA-derived sequences, their capacity to induce significant transcriptional activity through these repeats varies significantly (Gangwal et al. 2010). The FET family proteins FUS, EWSR1, and TAF15 are characterized by intrinsically disordered, low-complexity prion-like domains (PrLDs). These domains facilitate multimerization, physiological liquid-liquid phase separation (LLPS), and, under certain conditions, pathological protein aggregation (Thomsen et al. 2013; March et al. 2016; Ahmed et al. 2021).

In Ewing sarcoma, the multimerization of the EWS::FLI1 fusion protein via the EWSR1-derived N-terminal PrLD plays a pivotal role in recruiting the BAF complex to GGAAµSats (Boulay et al. 2017; Johnson et al. 2017). Similarly, the endogenous EWSR1 protein interacts with the BRG1 (SMARCA4) subunit of the BAF complex through its own N-terminal PrLD—a process that is likewise essential for the oncogenic function of EWS::FLI1 (Boulay et al. 2017). This interaction is disrupted by tyrosine-to-serine mutations within the EWS::FLI1 PrLD, suggesting that these proteins colocalize and function through a mechanism of co-phase separation (Ahmed et al. 2021).

Despite their prominent relevance in Ewing sarcoma, emerging evidence suggests that non-fused, full-length ETS transcription factors can also engage and activate GGAAµSats in other malignancies. For instance, in prostate cancer, the oncogenic ETS proteins ERG, ETV1, ETV4, and ETV5 have been shown to activate GGAA repeat reporters (Kedage et al. 2016). These oncogenic ETS proteins are overexpressed as full-length or N-terminally truncated forms in approximately half of all cases (Clark et al. 2007; Kedage et al. 2016). Furthermore, in B-cell acute lymphoblastic leukemia (B-ALL), ERG directly binds to and transforms GGAAµSats into active enhancers, thereby sustaining the leukemic gene expression signature (Kodgule et al. 2023). Crucially, this regulatory transformation is gated by the status of the ETS repressor ETV6: it occurs only upon mutations that cause the loss of ETV6 or the inactivation of its DNA-binding domain (Bohlander 2005). Under normal conditions, ETV6 binds these GGAA repeats to antagonize the activating effects of ERG, serving as a molecular safeguard against leukemic reprogramming (Kodgule et al. 2023).

Similar to FET::ETS fusion proteins, activation of GGAA microsatellites by full-length, aberrantly expressed ETS factors in prostate cancer is also suspected to require PrLD-dependent multimerization via FET family proteins (Kedage et al. 2016). However, in this context, in contrast to a chimeric fusion, multimerization is likely mediated via non-covalent protein-protein interactions between the full-length ETS factors and FET proteins. While current evidence strongly implicates EWSR1 in this process, definitive proof remains elusive because the ubiquitous expression and functional redundancy of FUS, EWSR1, and TAF15 complicate efforts to isolate their individual contributions within human cancer cells (Andersson et al. 2008).

We have previously shown that human EWS::FLI1 reproduces its neomorphic activation of transcription from synthetic GGAAµSat reporters in Drosophila (Molnar et al. 2022). Prompted by the evidence that full-length ETS factors can also activate these elements in human cancer cells, we aimed to explore the requirements for this activation using our Drosophila platform.

In this study, we systematically evaluate the transcriptional potential of various human oncogenic ETS factors at GGAA repeats and investigate the requirement for the single Drosophila FET orthologue, Caz. We show that ETV4 is uniquely capable of direct binding and activation of GGAA repeats in a length- and tissue-specific manner and demonstrate that the endogenous Caz protein is a critical determinant for converting these microsatellites into functional regulatory elements. We also show that human ETV6 functions as a strong inhibitor of this process in Drosophila. Moreover, by utilizing the TransTimer reporter system (He et al. 2019), we provide evidence of how human EWSR1 can substitute for its fly counterpart. Altogether, these findings provide substantial *in vivo* evidence of the functional conservation between human FET proteins and Drosophila Caz and confirm the essential role of endogenous FET protein in mediating the conversion of GGAAµSats into neoenhancers by non-fused oncogenic ETS transcription factors.

## RESULTS

### ETV4 Drives Length-Dependent Transcription from GGAA-Microsatellites

To evaluate the capacity of human ETS transcription factors to activate GGAA microsatellites (GGAAµSats) in Drosophila, we expressed human ERG, ETV1, FLI1, and ETV4 in various Drosophila tissues, including the salivary glands, accessory glands, and wing and eye-antennal imaginal discs. These experiments were conducted in a strain harboring the 20x(GGAA)µSat>TransTimer reporter, in which destabilized GFP (dGFP) reports recent transcriptional activity whereas the more stable RFP signal reflects cumulative transcriptional output over time.

While ERG, ETV1, and FLI1 failed to induce reporter expression in any examined tissue, ETV4 robustly activated the 20x(GGAA)µSat>TransTimer reporter in salivary and accessory glands, though not in imaginal or eye-antennal discs (Figure 1A; Supplemental Figure 1A). Supporting this observation, chromatin immunoprecipitation followed by quantitative PCR (ChIP-qPCR) from larval salivary glands expressing ETV4 (*hs-Gal4>UAS-ETV4*) demonstrated a specific enrichment of ETV4 at the 20xGGAA sequence (Figure 1B). These findings demonstrate that ETV4 is a potent activator of GGAA microsatellites in an *in vivo* context in Drosophila, a property not shared by the closely related ETS factors ERG, ETV1, or FLI1. Moreover, this capacity is strictly governed by the cellular context, as among the tissues that we tested activation was restricted to glandular tissues and absent in imaginal discs. This suggests that ETV4 requires a permissive environment—potentially involving specific co-factors or distinct chromatin states—to initiate transcription from these repetitive elements.

**Figure 1.**
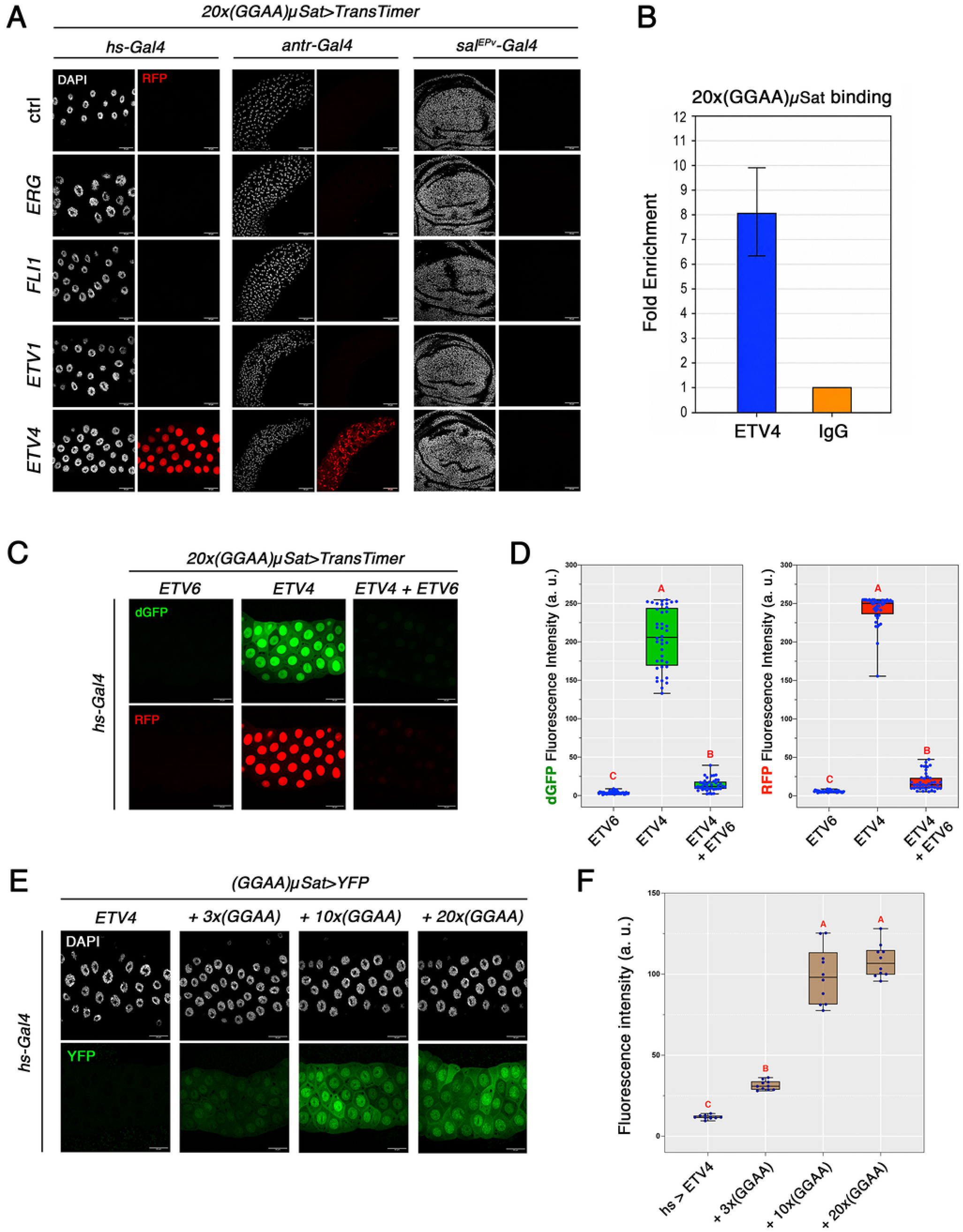
ETV4 drives tissue-specific and length-dependent transcription from GGAA-Microsatellites. (A) Representative confocal images of salivary glands, accessory glands and wing imaginal discs expressing human *ERG, ETV1, FLI1*, or *ETV4* under the control of *hs-Gal4, antr-Gal4* or *sal*^*EPv*^*-Gal4*. DAPI (grey) stained nuclei and RFP (red) reports transcriptional activation. Only ETV4 induces RFP expression from the 20x(GGAA)µSat reporter in salivary and accessory glands. Scale bar, 50 µm. (B) ChIP-qPCR showing enrichment of the 20x(GGAA) sequence in chromatin immunoprecipitated with anti-ETV4 antibody compared to IgG control. (C) Representative confocal images of salivary glands expressing ETV6 (*hs-Gal4>UAS-ETV6*), ETV4 (*hs-Gal4 > UAS-ETV4*), or both (*hs-Gal4 > UAS-ETV4; UAS-ETV6*). dGFP (green) and RFP (red) report transcriptional activation. ETV6 strongly reduces dGFP and RFP fluorescence. Scale bar, 50 µm. (D) Quantification of dGFP and RFP fluorescence intensity from glands shown in (B). Co-expression of ETV6 in ETV4 expressing glands strongly reduces both dGFO abd RFP levels. Different letters indicate statistically significant differences between groups (Kruskal–Wallis test followed by Dunn’s multiple comparisons test, adjusted *p* < 0.05). n = 40 nuclei per genotype (5 nuclei per salivary gland). a.u., arbitrary units. (E) Representative confocal images of larval salivary glands expressing ETV4 (*hs-Gal4 > UAS-ETV4*) and carrying (GGAA)µSat>YFP constructs with 3, 10, or 20 tandem GGAA repeats. DAPI staining (grey) marks nuclei and YFP (green) reports transcriptional activation. The 3x(GGAA)µSat reporter shows weak activation, whereas 10x(GGAA)µSat and 20x(GGAA)µSat reporters display strong YFP signal. All images were acquired under identical confocal settings. Scale bar, 50 µm. (F) Quantification of YFP fluorescence intensity in salivary glands for each reporter. ETV4-dependent activation was significantly lower from the 3x(GGAA)µSat reporter compared with the 10x and 20x reporters, indicating a repeat-length threshold for effective transcription. No significant difference was observed between the 10x and 20x reporters. Different letters indicate statistically significant differences between groups (Welch’S ANOVA followed by Dunett’s T3 multiple comparisons test, adjusted p < 0.05). n = 10 salivary glands per each condition. a.u., arbitrary units.

We then tested the effect in Drosophila of the ETS transcriptional repressor ETV6. In human B-ALL cancer cells, activation of GGAA repeats by oncogenic ETS proteins and the consequent dysregulation of the corresponding transcriptome signature can only occur upon inactivation of ETV6 (Bohlander 2005; Kodgule et al. 2023). Likewise, we found that expression of human ETV6 strongly suppressed ETV4-dependent transcription activation from 20x(GGAA)µSat down to background levels in salivary glands (Figure 1C,D). This inhibitory effect of ETV6 further underscores the parallelism between ETV4-driven activation of GGAA repeats in Drosophila and human cells. It also reinforces the idea that GGAA repeat activity arises from a dynamic interplay between activating and repressive ETS factors (Lu et al. 2023).

In human cells, the number of GGAA repeats is critical for ETS-driven transcriptional activation (Kedage et al. 2016; Kodgule et al. 2023). To determine the relationship between repeat length and ETV4-mediated activation, we quantified YFP fluorescence in salivary glands expressing ETV4 and carrying 3x, 10x, or 20x(GGAA)µSat>YFP constructs (Figure 1E,F). Compared to control flies lacking the reporter, the 3x(GGAA)µSat>YFP construct exhibited a modest but significant 3-fold increase in fluorescence (*adj. p* = 7.81 x 10^-10^). In contrast, both the 10x and 20x reporters showed a robust, 10-fold increase (*adj. p* = 4.35 x 10^-7^ and *adj. p* = 1.54 x 10^-9^, respectively). Notably, the levels of YFP upregulation elicited by ETV4 did not differ significantly between the 10x and 20x reporters (*adj. p* = 0.6547), suggesting a potential saturation of transcriptional activation at ten GGAA repeats. These results demonstrate that transcriptional output correlates with GGAA repeat length, reaching a plateau at ten repeats, consistent with cooperative binding models for tandem motifs (Guillon et al. 2009). Notably, this trend mirrors the EWS::FLI1-mediated activation of GGAA microsatellites observed in Drosophila, though the fusion oncogene requires a longer tract—twenty repeats—to reach saturation (Molnar et al. 2022).

### The single Drosophila FET orthologue Cabeza Regulates ETV4-Mediated GGAA-Microsatellite Transcription

In prostate cancer cells, EWSR1 is hypothesized to be an essential cofactor for oncogenic ETS factors to activate transcription from GGAA microsatellites (Kedage et al. 2016). However, evidence remains inconclusive because the effect of EWSR1 depletion is only partial, probably owed to the redundant functions of all three FET family members—FUS, EWSR1, and TAF15—which are ubiquitously expressed (Andersson et al. 2008).

Taking advantage of the fact that Drosophila possesses a single FET ortholog, Cabeza (Caz), we investigated whether Caz is required for ETV4-mediated transcriptional activity. To this end, we quantified the effect of Caz depletion on ETV4-driven transcriptional activation from the 20x(GGAA)µSat>TransTimer reporter (Figure 2A,B,C,D). Compared to a control RNAi (*mCherry*^*35785*^), RNAi-mediated knockdown of Caz using two independent lines (*caz*^*33990*^ and *caz*^*34839*^) in ETV4-expressing salivary glands resulted in a robust reduction of both the short-lived dGFP (*adj*.*p* = 1.16 x 10^-19^ and *adj*.*p* = 1.95 x 10^-33^, respectively) and the long-lived RFP fluorescence (*adj*.*p* = 1.1 x 10^-15^ and *adj*.*p* = 1.67 x 10^-20^, respectively) (Figure 2A,B,C,D). Given that the reported knockdown efficiencies for these lines are only partial, about 80% for *caz*^*34839*^ and 50% for *caz*^*33990*^ (Frickenhaus et al. 2015), these strong phenotypes indicate that Caz is an essential cofactor, necessary for ETV4-mediated transcriptional activation at GGAAµSats in the Drosophila salivary gland.

**Figure 2.**
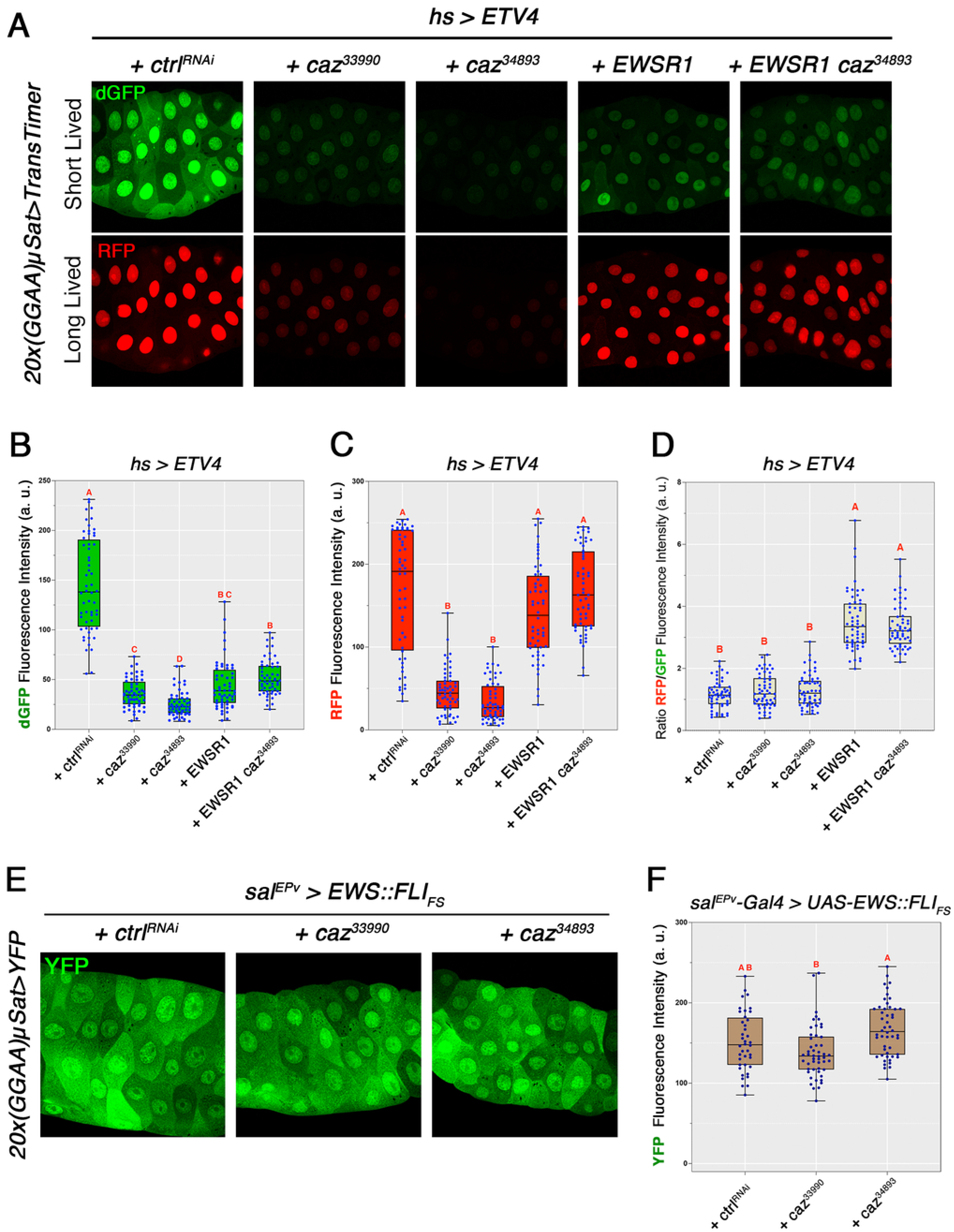
The Drosophila EWSR1 ortholog Cabeza is required for ETV4 to activate transcription from 20x(GGAA)µSat. (A) Representative confocal images of salivary glands expressing *hs-Gal4>UAS-ETV4* together with 20x(GGAA)µSat>TransTimer and either a control RNAi line (*ctrl*^*RNAi*^ *= mCherry*^*35785*^*-RNAi*), two independent *caz* RNAi lines (*caz*^*33990*^ or *caz*^*34839*^), human EWSR1, or human EWSR1 together with *caz*^*34839*^. Depletion of *caz* with either RNAi lines strongly reduced both short-lived dGFP and long-lived RFP fluorescence. Co-expression of EWSR1 in *caz*-depleted salivary glands (*EWSR1 caz*^*34839*^) partially restored reporter activity, with a stronger recovery of the RFP signal. (B-D) Quantification of dGFP fluorescence intensity (B), RFP fluorescence intensity (C) and RFP/dGFP ratio (D) from glands shown in (A). Both *caz* RNAi lines significantly reduced dGFP and RFP fluorescence. Co-expression of EWSR1 in *caz*-depleted background restored RFP levels to values comparable to those observed with EWSR1 expression alone. Depletion of *caz* did not significantly alter the RFP/dGFP ratio compared to the control, whereas co-expression of EWSR1 either alone or in the *caz*-depleted background, significantly increases this ratio. (E) Representative confocal images of salivary glands expressing EWS::FLI1 together with the 20x(GGAA)µSat>YFP reporter either *ctrl*^*RNAi*^, *caz*^*33990*^ or *caz*^*34839*^. (F) Quantification of YFP fluorescence intensity from the genotypes shown in (E). Depletion of *caz* did not cause a consistent reduction in EWS::FLI1-driven reporter activity. Groups not sharing a letter are significantly different from each other, as determined by Kruskal–Wallis test followed by Dunn’s multiple comparisons test, adjusted *p* < 0.05. For B–D, n = 56 nuclei per genotype (4 nuclei per salivary gland). For F, n = 40 nuclei per genotype (5 nuclei per salivary gland). a.u., arbitrary units.

To determine if the role of Caz in this context is functionally equivalent to human EWSR1, we quantified the effect of expressing human *EWSR1* on ETV4-driven transcriptional activation (Figure 2A,B,C,D). We found a significant difference between the short (dGFP) and the long-lived (RFP) reporters. In Caz wild-type salivary glands, *EWSR1* expression had no significant effect on the expression of the long-lived RFP signal, but significantly reduced the expression of dGFP fluorescence compared to the control RNAi (*adj*.*p > 0*.*9999* and *adj*.*p = 2*.*76 x 10*^*-15*^, respectively). However, more importantly, *EWSR1* expression in Caz-depleted cells (*caz*^*34839*^*-RNAi*) fully rescued both the RFP and dGFP signals, back to the levels observed in Caz wild-type salivary glands expressing EWSR1 (Figure 2A,B,C,D). These results demonstrate EWSR1 and Caz functional homology in the activation of transcription from 20x(GGAA)µSat by ETV4 in Drosophila salivary glands. In addition, the complete restoration of the cumulative RFP signal indicates that EWSR1 is fully capable of sustaining total transcriptional output over time. Nonetheless, the partial rescue of the short-lived dGFP signal—compared to endogenous levels—suggests that while EWSR1 supports the initiation of new transcription, it may possess lower kinetic efficiency than the native Caz within the Drosophila cellular environment.

Interestingly, ERG, ETV1, and FLI1, which failed to activate transcription for the GGAAµSat reporter when expressed on their own, remained silent upon coexpression with EWSR1 in salivary glands, accessory glands or wing discs (Supplemental Figure 2). This demonstrates that, like the endogenous Caz, human EWSR1 is not sufficient to confer transcriptional activity to any of these ETS proteins at GGAA repeats.

Mechanistically, our results are consistent with the proposal that the interaction between full-length EWSR1 and ETS proteins in human cancer cells mimics the activity of the corresponding oncogenic fusion which harbors its own PrLD and DNA binding domain. Consistent with this hypothesis, we found that YFP levels in Caz depleted cells in EWS::FLI1 expressing salivary glands remained indistinguishable from the control condition (*adj*.*p* = 0.1764 and *adj*.*p* = 0.124 for *caz*^*33990*^ and *caz*^*34839*^, respectively) (Figure 2E,F) indicating that the fusion oncogenic protein does not require Caz for GGAAµSat activation.

## Discussion

While the conversion of transcriptionally silent GGAA microsatellites (GGAAμSats) into functional enhancers is widely recognized as a hallmark driving the pathogenesis of Ewing sarcoma via FET::ETS fusion proteins like EWS::FLI1, this regulatory mechanism is also relevant across other cancer types through non-fused, full-length ETS transcription factors. In prostate cancer, overexpressed oncogenic ETS proteins such as ERG, ETV1, ETV4, and ETV5 actively engage and activate these GGAA repeat regions (Kedage et al. 2016). Similarly, in B-cell acute lymphoblastic leukemia (B-ALL), ERG transforms these microsatellites into active enhancers to sustain leukemic gene expression following the loss or inactivation of the ETS repressor ETV6 (Kodgule et al. 2023).

Using transgenic Drosophila lines carrying fluorescent reporters of GGAAμSat transcriptional activation, we have found that human ETV4 robustly activates transcription from GGAA microsatellites *in vivo* in the salivary and accessory glands, while ERG, ETV1, and FLI1 remain transcriptionally silent. Furthermore, ETV4 activity is highly context-dependent, as it does not activate GGAAμSat in other tissues like the wing and eye-antennal imaginal discs. These stark differences highlight a pronounced functional divergence among ETS factors with regards to their effect on GGAA repeats and reveal that tissue-dependent interactions with the Drosophila proteome are strictly required to engage and unlock these silent genomic elements.

Our study identifies Caz, the Drosophila orthologue of the human FET proteins as a critical requirement for ETV4 to activate transcription from GGAAμSats. Moreover, it also shows that human EWSR1 can partially substitute and compensate for the loss of Caz in the activation of transcription from 20xGGAAμSats by ETV4. These results demonstrate functional conservation between the fly and its human orthologue. Nonetheless, the observations derived from the GGAAμSat TransTimer reporter revealed a subtle difference in the kinetic profile for this rescue: human EWSR1 fully restores the cumulative transcriptional pool over time but only partially restores the “fresh” nascent signal compared to endogenous Caz. This suggests that while the human protein is functionally compatible, it may be less kinetically efficient within the Drosophila cellular environment than the native Caz ortholog.

Importantly, our finding that ERG, ETV1, and FLI1 remain unable to erase transcriptional silencing at GGAAμSats even when co-expressed with human EWSR1 further underpins the view that neither EWSR1 nor Caz is sufficient to confer microsatellite activation potential to all full-length ETS factors *in vivo*. This highlights the critical importance of additional lineage-specific cofactors, unique protein-protein interactions, or distinct differences in the baseline chromatin accessibility landscapes between mammalian and insect systems.

In prostate cancer, oncogenic ETS-dependent transcriptional activation of GGAAμSats has been suggested to be mediated through interaction with EWSR1 (Kedage et al. 2016). This observation led to propose that whether covalently tethered in a fusion protein or brought together through protein-protein interaction, the conjunction of an ETS DNA-binding-domain and a FET protein prion-like low-complexity domain can erase transcription silencing at GGAAμSats. Our results fully substantiate this hypothesis.

However, while sharing foundational similarities, the transactivation mechanisms driven by covalent tethering versus physical interaction exhibit three distinct differences. Firstly, while full-length FLI1 has no effect, the EWS::FLI1 fusion protein drives strong Caz-independent GGAAμSats transcriptional activation. Secondly, our data reveal that the transcriptional output of the full-length ETV4/Caz cooperative complex plateaus at ten repeats. In contrast, the EWS::FLI1 fusion oncogene requires a twenty-repeat saturation point in Drosophila (Molnar et al. 2022). This divergent length-dependence likely reflects differential cooperative binding of transcription factors and cofactors across multiple tandem motifs, which ultimately shapes chromatin accessibility and establishes distinct thresholds for effective enhancer output. Finally, the mechanisms driven by covalent tethering and physical interaction also differ regarding ETV6-mediated repression: in Drosophila salivary glands, ETV6 efficiently represses the activity of full-length ETV4 but has no effect on the EWS::FLI1.

Taken together, our findings support a model in which full-length ETS transcription factors can activate transcription at GGAAμSats through interaction with FET family proteins. They also underscore the potential relevance of GGAAμSat-driven unscheduled transcription in other tumour types beyond Ewing sarcoma.

Together, these findings demonstrate that GGAAμSats activation is a broader oncogenic strategy shared by various malignancies beyond its classic context in Ewing sarcoma.

## Acknowledgements

We are very grateful to S. Llamazares; Bloomington Drosophila Stock Center; and to all members of our laboratories for very helpful discussions. We are also grateful to the Band of Parents at Hospital Sant Joan de Déu for supporting the overall research activities of the Developmental Tumor Laboratory, Pediatric Cancer Center Barcelona (PCCB).

## Data Availability

Strains and plasmids are available upon request. The authors affirm that all data necessary for confirming the conclusions of the article are present within the article and figures.

## FIGURE LEGENDS

Figure S1. ETV4 does not activate 20x(GGAA)µSat>TransTimer in eye discs.

(A) Eye imaginal discs from larvae expressing mCD8-GFP or ETV4 in cells posterior to the morphogenetic furrow with longGMR-Gal4. ETV4 expression has no effect in the ommatidia development.

Figure S2. Human EWSR1 has no effect on ETS factors to activate expression from 20x(GGAA)µSat in Drosophila.

Representative confocal images of salivary glands (A), accessory glands (B) and wing imaginal discs (C) co-expressing human ERG, ETV1, FLI1, or ETV4 with EWSR1 under the control of *hs-Gal4, antr-Gal4* or *sal*^*EPv*^*-Gal4*. DAPI (grey) stained nuclei and RFP (red) reports transcriptional activation. Only co-expression of EWSR1 and ETV4 induces RFP expression from the 20x(GGAA)µSat reporter in salivary and accessory glands. Scale bar, 50 µm.

## METHODS

### Fly stocks

The following fly stocks were used in this study: *antr-Gal4* (Lee et al. 2018), *sal*^*EPv*^*-*Gal (Cruz et al. 2009), *hs-Gal4* (Bloomington Drosophila Stock Center (BDSC) #1799), *longGMR-Gal4* (BDSC #8121), *UAS-mCD8-GFP* (BDSC #5130), *UAS-EWSR1* (BDSC #79592), *UAS-caz*^*RNAi*^ (BSDC #32990 and #34839), and *UAS-mCherry*^*RNAi*^ (BSDC #35785). The GGAAµSat reporter lines 20x(GGAA)µSat>YFP, 10x(GGAA)µSat>YFP, and 3x(GGAA)µSat>YFP, as well as the *UAS-EWS::FLI1*_*FS*_ line and *UAS-FLI1*, were previously described (Molnar et al. 2022). The following transgenic lines were generated and used in this work: *UAS-ERG, UAS-ETV1, UAS-ETV4, UAS-ETV6*, and *20x(GGAA)*µ*Sat>TransTimer*. All crosses, including controls, were maintained at 25ºC.

### Immunohistochemistry and microscopy

Immunostaining of salivary glands, wing discs, and accessory glands was performed as follows. Salivary glands and wing discs were dissected from wandering third instar larvae, and accessory glands from 4-5 day-old adult males, in phosphate-buffered saline (PBS). Tissues were fixed for 30 min in 4% formaldehyde with 0.3% TritonX-100, followed by three washes of 15 min each in PBS with 0.3% Triton X-100 (PBST). DNA was stained with 4’,6-diamidino-2-phenylindole (DAPI). Salivary glands, wing discs and accessory glands were mounted in Vectashield (Molecular Probes). Images were acquired with a Leica SP8 confocal microscope. The objective used was HC PL APO CS2 40x/1.30 OIL. Images were acquired at zoom 1. All immunofluorescence images shown correspond to a single optical section (Z-plane) and were processed using ImageJ.

### Quantification and statistical analysis

Fluorescence quantification in salivary glands was performed using ImageJ by calculating mean grey values in a single focal plane per gland from images acquired with a Leica SP8 confocal microscope. For the 3x, 10x, and 20x(GGAA)µSat>YFP reporter lines, mean grey values were measured for each region of interest (ROI) per salivary gland (10 salivary glands per genotype). For the 20x(GGAA)µSat>TransTimer reporter line, mean grey values of dGFP and RFP fluorescence were quantified in four nuclei per salivary gland (14 salivary glands per genotype). All individual measurements were represented as box plots, and statistical analyses, including *p*-value calculation were performed using GraphPad Prism 10 for MacOS. Statistical significance was assessed using Welch’s ANOVA followed by Dunnett’s T3 multiple comparisons test for YFP measurements, and using the Kruskal-Wallis test followed by Dunn’s multiple comparisons test for dGFP and RFP measurements. Compact Letter display (CLD) was used to indicate statistically significant differences in box plots.

### Cloning and transgenesis

Sequences codon optimized for *Drosophila melanogaster* corresponding to the proteins ETV4 (P43268-1), ETV1(NP_001357485.1), and ERG (P11308-4) were synthetized by GeneWiz. ETV6 (P41212) ORF was synthetized by Twist Bioscience. All the ORF was cloned in the *pUASt-attB* vector in *EcoRI*-*XbaI*. The 20x(GGAA)µSat>TransTimer reporter was made by inserting a PCR fragment containing the transcriptional timer (He et al. 2019) in the *p20x(GGAA)*µ*Sat-YFP* vector (Molnar et al. 2022) in *EcoRI-BglII*. The fragments were amplified from *pUAST-IVS-syn21-nlsGFP-PEST-2A-nlsRFP-p10* (DGRC Stock 1498). The primers used were: *UP-ATG*: 5′-AGGGAATTGGGAATTCCAAAATGCCCAAGAAGAAGC-3′; *UP-PEST*: 5′-GCTTCTGCTAGGATCAATGTG-3′; *DO-PEST*: 5′-CACATTGATCCTAGCAGAAGC-3′ and *DO-STOP*: 5′-CCGCGGCCGCAGATCTTTATTTTAAAAACGATTCATTCTA-3′.

All PCR amplifications were performed using KOD polymerase (Merck). Cloning was performed using In-Fusion Snap Assembly (Takara). Primers were ordered from Sigma.

Transgenic fly lines were generated by BestGene Inc. using the attP16 docking site for the 20x(GGAA)µSAt>TransTimer transgene, the *M{3xP3-RFP*.*attP’}ZH-51C* docking site for the *UAS-ETV1, UAS-ERG*, and *UAS-ETV4* transgenes, and the *M{3xP3-RFP*.*attP}ZH-86Fb* docking site for the *UAS-ETV6* transgene. All transgenic lines used in this study were sequenced to confirm the presence and integrity of the correct transgene.

### ChIP-qPCR

ChIP experiments were performed essentially as described (Climent-Cantó et al. 2020). A total of 300 larval salivary glands were dissected in 1xPBS and fixed with 1.8% formaldehyde for 10 min at room temperature. Samples were then washed in wash buffer A (10mM HEPES pH 7.9, 10mM EDTA, 0.5mM EGTA, 0.25% Triton X-100), followed by wash buffer B (10mM HEPES pH 7.9, 100mM NaCl, 1mM EDTA, 0.5mM EGTA, 0.01% Triton X-100). Salivary glands were pelleted, resuspended in 900 µl of lysis buffer (10mM Tris-HCl pH 8.0, 1mM EDTA, 1% SDS and 1mM AEBSF), and sonicated using a Bioruptor Pico sonication devide (Diagenode) for nine cycles of 30 s ON and 30 s OFF to obtain chromatin fragments of 200-500 bp. After sonication, lysates were adjusted to 1% Triton X-100, 0.1% sodium deoxycholate (DOC), 140mM NaCl, 1% SDS and 1mM AEBSF. Samples were centrifuged and the supernatant was collected. For each immunoprecipitation, 1 ml of chromatin was used, and 40 μl were saved as input sample. Chromatin aliquots were pre-cleared with 50 µl of Dynabeads Protein A (Thermo Fisher Scientific) for 3 h at 4 °C. Immunoprecipitations were performed using anti-ETV4 antibody (E1W1G, Cell Signaling Technology) or anti-DYKDDDDK Tag antibody as mock control (D6W5B, Cell Signaling Technology). A total of 4 µg of the corresponding antibody was added to each sample and incubated overnight at 4°C on a rotating wheel. The following morning, 50 µl of BSA-blocked Dynabeads Protein A were added to each sample and incubated for 3h at 4 °C. Beads were then collected on a magnetic rack and washed with RIPA buffer, followed by one wash with LiCl wash buffer (250mM LiCl, 10mM Tris-HCl pH 8.0, 1mM EDTA, 0.5% NP-40, 0.5% DOC) and two washes with TE buffer. Beads were treated with DNase-free RNase A/T1 (Thermo Fisher Scientific) for 30 min at 37°C. Crosslinks were reversed by adjusting samples to 1% SDS, 0.1M NaHCO_3_ and 0.2 mg/mL Proteinase K, followed by overnight incubation at 65°C. DNA was purified using the QIAquick PCR Purification Kit (QIAGEN), according to the manufacturer’s instructions.

Purified DNA from two independent ChIP experiments was analysed by quantitative PCR using PowerUp SYBR Green Master Mix (Thermo Fisher Scientific). qPCR was performed using primers targeting the 20x(GGAA)µSat region and an SV40 region of the reporter construct as a control region. The primers used were: *20xGGAA-+1F*: 5’-CATGCCTGCAGGTGAGGGAG-3’; *20XGGAA-R*: 5’-CTCGCTAGAGTCTCCGTTTCTTTTCC-3’; *SV40-F*: 5’-TGCTGACTCTCAACATTCTACTC-3’; and *SV40-R*: 5’-CTCTAGTCAAGGCACTATACATCAA-3’. The qPCR results were analyzed using the fold change method, and ChIP enrichment was calculated relative to input DNA.

